# Investigation of a mathematical model describing global cancer growth and treatment: An inhomogeneous model based on the generalized logistic equation

**DOI:** 10.1101/2022.12.18.520960

**Authors:** Haofan Wang, Yitao Mao, Zhen Zhang, Zhenxiong Xu, Shuyang Luo, Weifeng Li, Sibin Zou, Bin Chen, Huiquan Wen, Longxin Lin, Weihua Liao, Mingsheng Huang

## Abstract

Tumor growth is manifestation of the evolution of a complex system. Researchers have limited scope of modelling studies on specific aspects or stages of the process. It has led to unsatisfactory explanation of clinical observations. We hereby demonstrated that an inhomogeneous model built on the generalized logistic equation could serve better. It was developed to describe the whole process of tumor progression, clinically observed independence of index tumor growth from spread of the disease and growth deceleration during early stage of solid tumors. It was validated by simulating the coexistence of exponential and sigmoidal growth in chronic lymphocytic leukaemia, theories of tumor heterogeneity, as well as by accommodating notions pertaining to tumor treatment and prognosis. We thought therefore it was an interesting and not unjustifiable description of actual tumor growth in human body and hoped it might encourage more researchers to look at tumor modelling from a clinical perspective.

## Introduction

Mathematical simulation of expansion of all cancer cells in a patient from early to advanced stage is important both in a clinical sense and for researchers. Standard models have been helpful but also shown insufficiencies describing this complex process. Standard exponential growth has been considered an essential feature of malignant tumor^1–17^. It provided mathematical support for selecting 5 years as the standard of cure^1, 14, 16–19^. Many researchers considered exponential function an A priori hypothesis for modelling studies^1–4, 9, 13, 15^. However, deceleration in growth or significantly slower growth of single large tumors compared to smaller ones has long been recorded^2, 10, 16–32^. Any slowdown is against exponential growth. Biologically, exponential growth would require cancer cells to grow completely freely in a patient^1^ but this is impossible because at the very least the carrying capacity of human body would pose an ultimate restriction.

Equations that allow for limiting factors including Logistic, Gompertz, Bertalanffy and power-law equations have been studied^4–12^. However, explaining the above-mentioned deceleration by these equations was attached with a rather awkward prediction that these tumors should stop growing before they reached clinically advanced stage^17, 20–31^. In hepatocellular carcinoma (HCC), it has been demonstrated that assuming a reported cell-doubling time of 204±132 days, Gompertzian growth would allow HCCs to grow only to around 10-12cm^11^. A solitary HCC around 10cm in diameter in current guidelines is often categorized as early stage^19^. Studies showing better identifiability of sub-exponential models have acquired tumor growth data from singular nodules or benign lesions ^10, 11, 17, 20–36^, indicating that these models fitted only localized growth upon which local and thus most likely physical restrictions apply. This would lead to insufficiency though, should the model be required to also approximate growth of the whole cancer cell population in advanced stage malignancy, because an essential feature of malignant tumor as opposed to benign tumor was that invasive and metastatic growth could overcome limitations on primary lesions at this stage. Clinical evidence suggest growth acceleration could happen in advanced stage cancers ^3, 19, 41, 42^. For example, portal vein tumor thrombosis was once demonstrated to grow faster when it was larger ^41^. Acceleration after stalled growth has also been observed clinically in liver cancer lesions ^21, 24, 25, 31^. Mathematically, models allowing free cell movement across local boundary predicted exponential growth ^1, 37, 38^. It is also clear that if all lesions are growing according to one Gompertzian equation, increase in total cancer cell population should not be represented by a similar model, especially considering that primary and secondary seeding from all colonies are generating new lesions at the same time ^3, 9, 12, 39, 40^. Up till now, there has been no discussion to our knowledge of a model describing growth of the whole cancer cell population that covers such transition from sigmoidal growth under local restrictions at early stage to accelerated progression beyond these boundaries at advanced stage.

Differential equations of tumor growth also predict evolution of genetic heterogeneity. Exponential models command a tendency towards homogeneity^1, 12, 13, 43^. Suppose we have one clone of cells growing at rate a1 and a faster one a2. Then

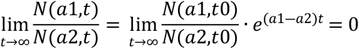

where N(t) denotes the number of tumor cells ^43^. Biologically, mobility is essential for tumor cells to fulfill exponential potentiality because it provides means for faster growing clones to overrun others, also known as the ‘sweeping model’ ^1, 13, 37, 38^. Clinico-pathological evidence though is inconsistent regarding homogeneity ^44–54^. Evidence of extremely high clonal diversity in large solid tumor has been exhibited ^53^. On the other hand, although many supported that HCCs are heterogenous, a general correlation between larger tumors and worse differentiation was also recognized^13, 44–46^ which suggested sweeping like mechanism. Insufficient replacement suggested a more complicated growth model than described by standard equations.

Clinical implication of tumor growth models was built on the assumption that increase in size and number of tumor lesions was driven by increase in number of tumor cells. This went in line with size and number of tumor lesions being the most prominent prognostic factors when we computed and compared predictive values of multiple baseline variables by projecting them onto survival time vector or when we set up treatment guidelines ^19, 55^. The question was though, if both size and spread were propelled by one differential equation of tumor growth, they should not have been separated into independent factors. Yet number and size of lesions have constantly been tabulated as independently significant prognostic factors ^19, 55^, sometimes even the only two that survived after computational analysis had clipped dozens of other candidates based on data from large group of patients ^55^. Pathological and modelling studies provided more direct evidence that invasion and metastases developed independently from tumor growth at site of origin ^2, 18, 56^. Standard models would not allow this independence of growth of primary tumor from spread of the disease beyond local restrictions. Such independence has therefore not been reflected in metastasis models based on such standard tumor growth laws ^3, 9, 12, 38–40^.

These problems of standard models seemed to us lay in inferring growth models based on theories and experimental data ^1–13, 15^. Tumor development in vivo is affected by a network of molecular, cellular, local, and systemic processes. Growth models are intended to represent the composite effect of all these mechanisms. Laboratory experiments mostly had to control all other factors to illustrate one pathway. Data thus acquired are therefore not sufficient for Bayesian inferences. There are for example gaps among predictions of metastatic mechanism based on cytological data, xenograft models and clinico-pathological observations^3, 4, 9, 12, 48, 49^. This prompted us to look for a model describing growth of total tumor burden of any cancer throughout its lifespan from a clinical and phenomenological perspective.

## Methods

### Parameters set up

Tumor-growth simulation processes described below were carried out in Python (ver. 3.7, https://www.python.org/). Growth was described by the following equation (equation 1):

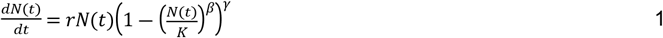

The generalized logistic equation was adopted to avoid making unnecessary assumptions of probability distribution by using standard forms of equations which assume exponents *β* and *γ* to be equaling 1 ^5, 43^. There should have been an exponent α for the first *N*(*t*) in the right-hand side (r.h.s) of equation 1 though. We assumed that it equaled 1 because we agreed that cancer cells would if given full support, proliferate exponentially^1–4, 6, 8, 9, 12, 48^. This oncological consideration was the main reason we did not use Bertalanffy or power-law equations. Parameter r represented birth to death ratio as in previous studies ^1, 14^. Larger r indicates potentiality of higher growth speed.

Parameter *β* described resistance to adverse environment since it appeared as exponent of 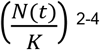^2–4, 6, 8, 9, 12, 48^. These two parameters represented inheritable features of cell clones and evolved with time. They provided means for describing heterogeneity in our model. Genetic and morphologic heterogeneity in various types of cancers from early to late stages have been demonstrated to correlate with various growth types and imaging features ^1, 12–14, 18, 19, 44, 45, 50–54^. These signs of heterogeneity were exhibited to accumulate and develop over time ^14, 44, 46, 50, 52, 54^. The widely accepted cancer stem cell and epithelial-mesenchymal transition theories supported this notion ^44^. Macroscopically, ‘nodule in nodule’ growth in solid tumors has been noted in different stages indicating emergence of faster growing and more resilient clones throughout the ‘life span’ of a tumor ^44^. Accordingly, we stipulated that r and *β* of any clone represented an average value that increased with mutational events. Therefore, two additional parameters were introduced to describe these alterations. which were:

Sr: the augmentation of r that a child clone get from its mother clone.
S_β_: the augmentation of β that a child clone get from its mother clone.

Parameter *K* in this study was set to be constant during each simulation, representing carrying capacity of individual patient. In future studies however it could variate with time or after treatment. Variable *γ* represented growth impeding effects other than carrying capacity. Both stochastic events and continuous influences participate in its evolution ^8, 51^. It was a time dependent variable described by:

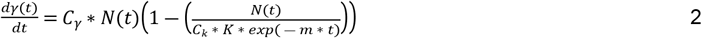

Three γ-related parameters were introduced. They were:

*C_γ_* represented growth restriction that increased with tumor size when it was in early stage ^8, 44, 51^. For example, it has been reported that when sectioned, HCC mass could “bulge above the cut surface” due to strangulation by pseudo-encapsulation ^51^. Computationally, a larger *C_γ_* led to faster growth of *γ*.
*C_k_* denoted innate invasive potential of tumor. It stemmed from the notion that HCCs switched to more aggressive growth over certain size^29^. Larger *C_k_* could translate to a later inflection point of the *γ* curve.
*C_k_* was multiplied by *e-^mt^* because we believed that invasive or metastatic colonies occurred randomly and would then continue to grow in an exponential-like way in their respective early stages of development^1–4, 8, 9, 12, 39^.

An additional parameter λ not shown in equations 1 & 2 described the probability of mutation each day. Thus, we defined eight parameters in our tumor growth simulation model: *m, C_γ_, C_k_, S*_r_, S_β_, λ, *r*, and *β*. In the simulation process, *r* and *β* were initiated by r_0_ and β_0_, indicating *r* and *β* of the very first clones. Initial value of *γ* was set to be 1.

### Tumor growth simulation

We used a recursion way to calculate the tumor cell number for each day by equation 1. Recursion step interval was set at one day. Supposed mutated cell number of each day was computed by N*λ. N represented population of each clone which was defined as all cells with same *r* and *β* on that day. And we assumed that mutation occurred randomly and independently. For example if clone A had N cells at the beginning of one day, and n cells out of N were supposed to be mutating, then the number of cells in A was reduced to (N-n). Meanwhile a new clone with n cells was produced. If was named clone A’, and if the *r* value and *β* value of clone A was r_0_ and β_0_, respectively, then r value and β value of A’ would be (1+S_r_)*r_0_ or (1+S_β_)*β_0_, because we stipulated that one mutational event would lead to only either change of r (r-type mutation) or *β* (β-type mutation), but not both at the same time. After the clones were updated, we calculated the number of cells for each clone on that specific day according to equation 1. Then the tumor cell number for each single clone and the total tumor cell number for the whole disease were updated accordingly. Simultaneously, *γ* would also be updated according to equation 2. Considering its biological implication, it was arbitrarily dictated that *γ* would not be less than 1. If the calculated *γ* on a specific day was less than 1, it would be replaced by 1. This process iterated day by day until the end point was reached. The end point was defined as the first day total cancer cell number reached *K* or surpassed *K*.

We assumed that the solid tumor was a standard spheroid, and that each single tumor cell had a diameter of 12×10^-4^ cm. Then the diameter of the whole solid tumor was derived from the following equation:

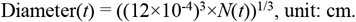

A curve of total disease volume versus days and one of the *γ* value versus days was generated by data from these processes. Curves describing volume of each clone versus days were also produced.

In the result section, we replaced the absolute diameters with their corresponding percentiles to transfer the data to a uniform range of 0-100%.

### Metastasis prediction

Researchers have investigated models based on standard equations for computation of probable start of metastasis using clinically acquired data of primary lesion growth and assumption of a universal seeding to original tumor size ratio ^12, 38–40^. Named by some as colonization rate, this ratio *β*(*x*) of a lesion with *x* cells is computed by ^40^:

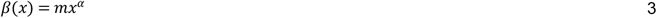

Parameter *α* in the r.h.s of equation 3 represented proportion of cells susceptible to yield metastasis in such tumor and *m* represented the per day per cell probability for such cells to overcome all the steps of the metastatic cascade. It is common among these studies to assume all lesions grow according to Gompertzian law^12, 39, 40^:

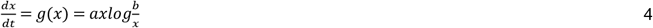

Conforming with the assumption that both primary and secondary seeding contributed to metastases our model would suggest that for computational simplicity, growth of all lesions (sources of seeding) combined be represented by standard exponential model ^1, 37^ instead of sigmoidal model. Using the same idea of a colonization rate *β*(*x*) that represented only distant metastasis, and assuming that each metastatic lesions grew according to one Gompertzian model, we would figure out the parameters by setting up a group of equations based on consecutive size measurements of the largest metastasis and the increment in number of observable metastases over time. Number of observable metastases *N\** that grew from day *t\** over a period of 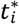 days can be calculated by:

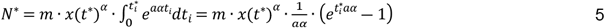

And number of cells in any one metastatic lesion *i* at time *t* can be expressed as:

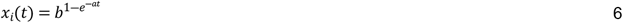

Reading figure 2 and figure 4a from the report by Iwata, K. and colleagues ^40^, it was determined that at the 432nd day after diagnosis, 10 metastases were found among which the largest one was computed to contain 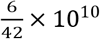 cells; 559 days after diagnosis, 18 more metastases were found with the largest one holding 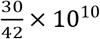 cells; and at the 632nd day, 48 metastases were found with the largest one holding 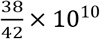 cells. Following equations were therefore constructed based on equations 5 and 6:

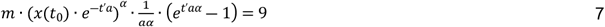

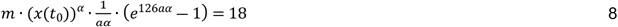

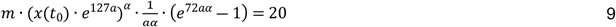

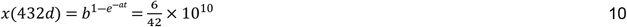

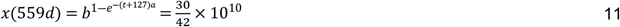

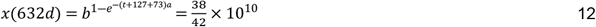

Variable *t’* represented time between inception of the 1^st^ and 11^th^ metastatic lesion while *t* in equation 10 represented time from inception of the first metastasis to its first diagnosis while *t*_0_ indicated the expected time of inception of the 11^th^ metastatic lesion.

#### Treatment simulation

For simulation of recurrence after surgery ^56^, we arbitrarily removed all but ten tumor cells representing surgical intervention. These were randomly picked up from all clones and grew de novo according to equation 1.

Systemic treatment was simulated by first-off removing a certain proportion of cells in each clone. We determined this proportion to be (*r* / *β*)%, i.e., after the treatment, the number of cells left in each clone would be [1 – (*r* / *β*)]% of pre-treatment population. We also simulated a recent theory of drug-tolerant persister state ^57^ by increasing the γ value during treatment. Treatment for one time as well as several times were simulated.

## Results

Growth curves generated by our model demonstrated that decelerated progression observed in early-stage solid tumors and reacceleration in advanced stage could be simulated (Figure 1a, see also Figure S1) ^19, 20–26^. Slowdowns that lasted for a short or lengthened period could happen either when tumor was small or when its size came near carrying capacity. Their occurrence was dictated by development of variable γ. It corresponded well with the definition of γ since early-stage was defined as tumor growth before significant breakthrough of regional limitations at site of origin which in our model was represented by before γ value was first computed to be less than 1.

**Figure 1.**
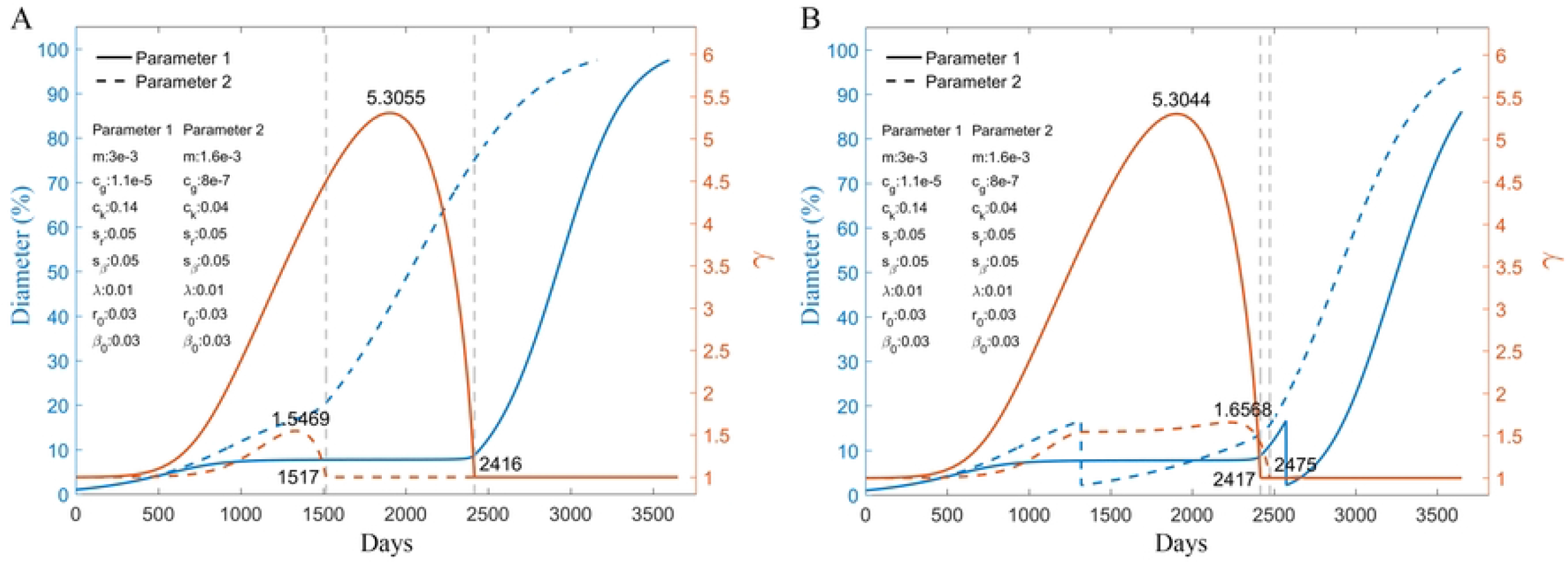
Modelling early-stage growth deceleration and multinodular tumor growth in relation to surgical management. A, Growth deceleration at early stage and acceleration later on was simulated. Curves that exhibited extended or short deceleration period when tumor size was relatively small were generated. These curves could explain reported different surgical outcome in seemingly similar early stage multinodular HCCs. We assumed multicentric occurrence (MO) type to have weaker local restriction (c=8·e^-7^) and lower breakthrough threshold (C_k_=0.04). Metastases formed in IM type when γ returned to 1 at around day 2416 (blue curve represented growth curve of IM type tumor while brown curve represented evolution of parameter γ of this type of growth with time). Assuming IM tumors were diagnosed at this point, MO type reached similar size at around day 1517 (blue dashed curve represented growth curve of MO type tumor while brown dashed curve represented evolution of parameter γ of MO type). B, Surgical intervention was simulated by randomly deleting all but 10 cells before γ of MO type (dashed curve) started to drop and when IM (blue curve) type was of similar size. It took MO type longer to regrow to size comparable to pre-surgical level. We interpreted it as compatible with the reported better recurrence free survival outcome of MO type cancer ^56^. Out of 100 trials, average time from surgery to 97.5% of K was 2688.06 ± 123.03 days for MO type and 1461.73 ± 154.91 days for IM type (data not shown in the line graph). Likewise, it corresponded to the reported longer overall survival of MO type ^56^. Similar assumption would also predict faster tumor growth at the time of recurrence, which was in accordance with accounts from an earlier clinicopathological study ^18^.

To demonstrate how our model would suggest a way of explaining clinical observations different from previous modelling studies based on standard equations, we extracted clinical measurement data from one of the papers that established a well acknowledged method of calculating metastatic probability for prediction of inception time of metastases based on standard Gompertzian model ^40^ and run prediction of metastasis forming time by our method. Computational results showed that

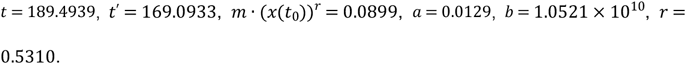

Variable *t’* represented expected time between inception of the 1^st^ and 11^th^ metastatic lesion while *t* in equation 10 – 12 represented expected time from inception of the first metastasis to its first diagnosis and *t*_0_ indicated the expected time of inception of the 11^th^ metastatic lesion. Colonization rate of primary and secondary seeding combined at *t*_0_ was described by *m* · (*x*(*t*_0_))*^r^*. Parameters *a* and *b* represented proliferative advantage and saturation level of the studied metastatic nodule, respectively.

Coexistence of exponential and sigmoidal growth as well as intermediate pattern observed in chronic lymphocytic leukaemia (CLL) ^14^ was also simulated (Figure 2, see also Figure S3). It validated the feasibility of our model because such was a particular phenomenon of natural cancer progression independent from that of varying growth speed in solid tumor. Noticeably, relation between faster mutation plus replacement when there was minimal local boundary effect and exponential growth also paralleled this and other reports ^1, 14, 37, 38^.

**Figure 2.**
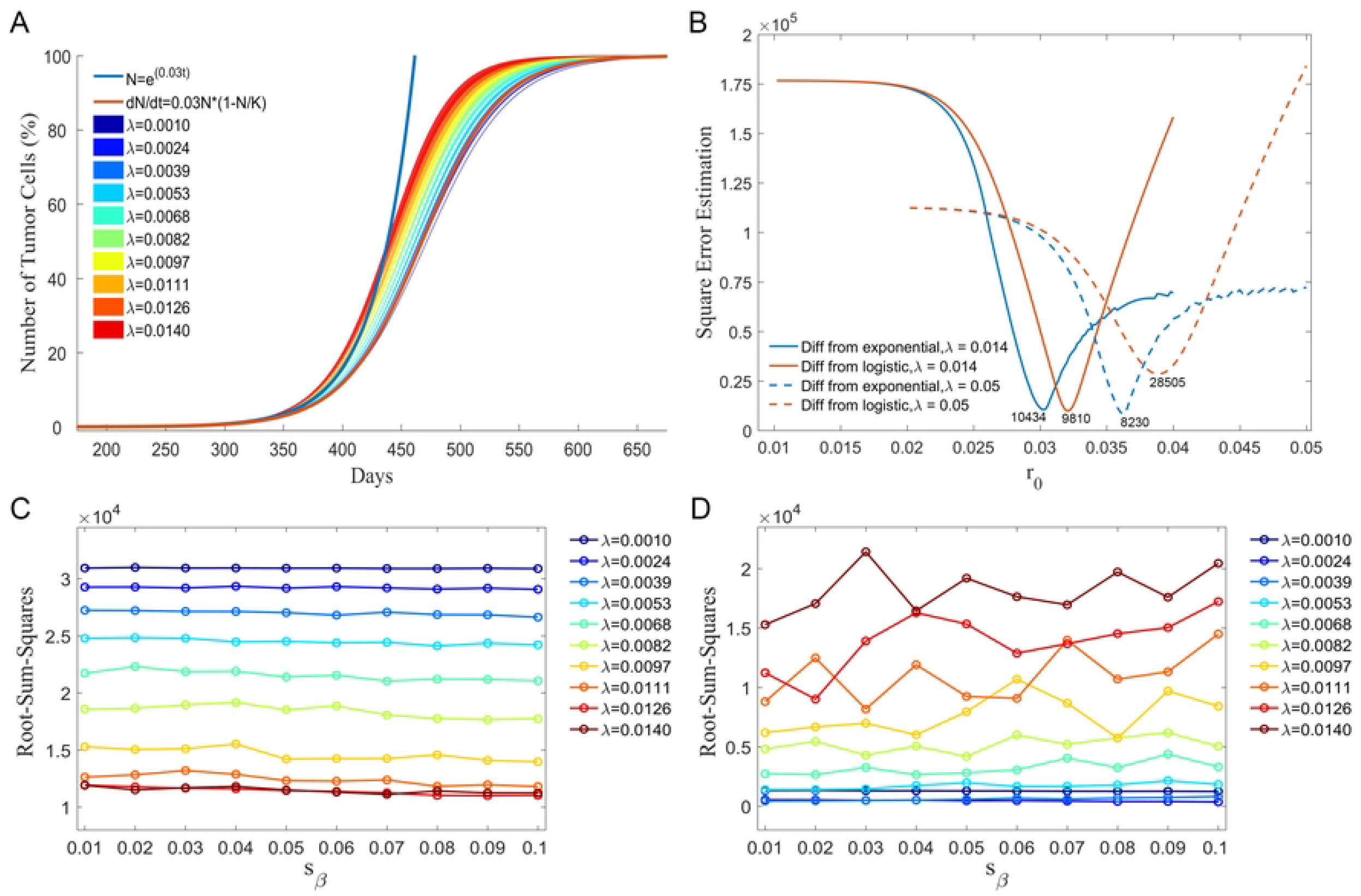
Simulation of coexistence of exponential like and logistic like growth in chronic lymphocytic leukaemia (CLL) A. Curves generated by our equation with higher rate of mutation denoted by λ were closer in shape to the exponential function with same parameter of birth-death ratio (r=0.03) than curves generated by the same equation with lower rates of mutation, which were closer to the logistic equation with same r. All curves in Figure 2A were given same value for all parameters except λ and S_β_ (m=5e^-5^, c_g_=e^-12^, C_k_=0.75, S_r_=0.06, r_0_=0.03, β_0_=0.9). We experimented with 10 different λs each marked in 2A with specific color. For each λ we tested 10 different values for S_β_ ranging from 0.01 to 0.1 at 0.01 intervals. The resultant curves were merged into respective colored bands. B. Experiment with our model (all other parameters remained unchanged while λ being set as 0.014 or 0.05), exponential functions (blue curve) and logistic curves (brown curve) with parameter r0 ranging from 0.01 to 0.04 (horizontal axis) showed that for the experimented model with 0.014 mutation rate, logistic curve would generate a closer approximation to our presumed natural tumor growth curve. Similar experiment with exponential functions (blue dashed curve) and logistic curves (brown dashed curve) with parameter r0 ranging from 0.03 to 0.05 would produce a best approximation that was closer to our tumor growth model with a relatively higher mutation rate (λ=0.05) than the best approximation from logistic model, which could explain why this type of tumor natural growth had been recognized as exponential in previous clinical reports^14^. C&D. When we took the summed value of squared differences (vertical axis) between two curves as a measure of approximation, data of these experiments showed that the exponential curve (C) was closer to a higher range of λ while the logistic curve (D) to a lower range of λ.

Also independently, it was recently suggested that all cancer cells from genetically different clones would enter a reversible drug-tolerant persister (DTP) state to evade death from systemic treatment ^58^. The researchers pointed out that this state was stochastically induced and preserved heterogeneity ^54, 58^. These observations could be simulated with our model by a sudden loss of cells in each clone proportional to its β value and concomitantly an arbitrary increase of γ (Figure 3).

**Figure 3.**
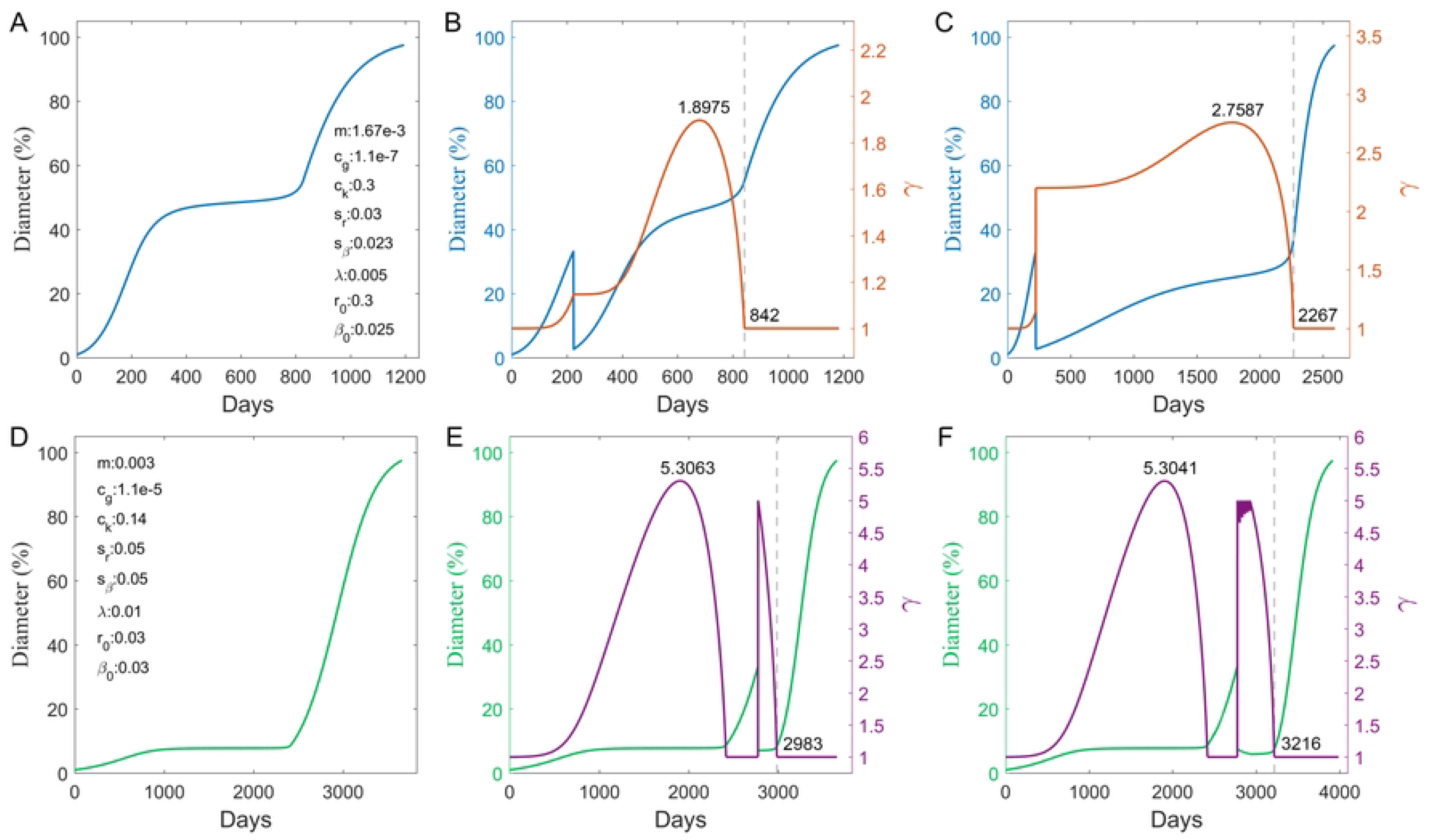
Treatment simulation. A. In this experiment natural growth of a tumor ended at 97.5% of K as surrogate for endpoint of total survival after 1318 days. B. This time was 1298 days if one treatment (Materials and Methods) without extra change in γ was given when the tumor was still confined by local restriction at site of origin. C. The same treatment was simulated again but with concurrent γ (brown curve) elevation to an arbitrary number 2.2 which was approximately the peak γ value of the tumor if left untreated (See also Figure S1A). The assumed overall survival was prolonged to 2689 days. D. Natural growth of another supposed tumor was simulated which would reach 97.5% of K in 3848 days. E. One session of γ elevating treatment was shown to bring overall survival under this condition to 3833 days if given after local restriction was breached. F. When same treatment was repeated 6 times at 30-day intervals, survival time was shown to be prolonged to 3864 days. Purple line depicted evolution of γ with time (gamma evolution of untreated tumor was shown in Figure 1).

Necrosis was recently shown to portend worse prognosis, which was somewhat predicted in an earlier modelling study 1, 59. In our model, necrosis was simulated not unlike treatment without concurrent increase in γ. It was shown to result in shortened survival (time to 97.5% K) and decreased peak γ level, which agreed with the reported correlation between necrosis and vascular invasion (Figure 4) 59.

**Figure 4.**
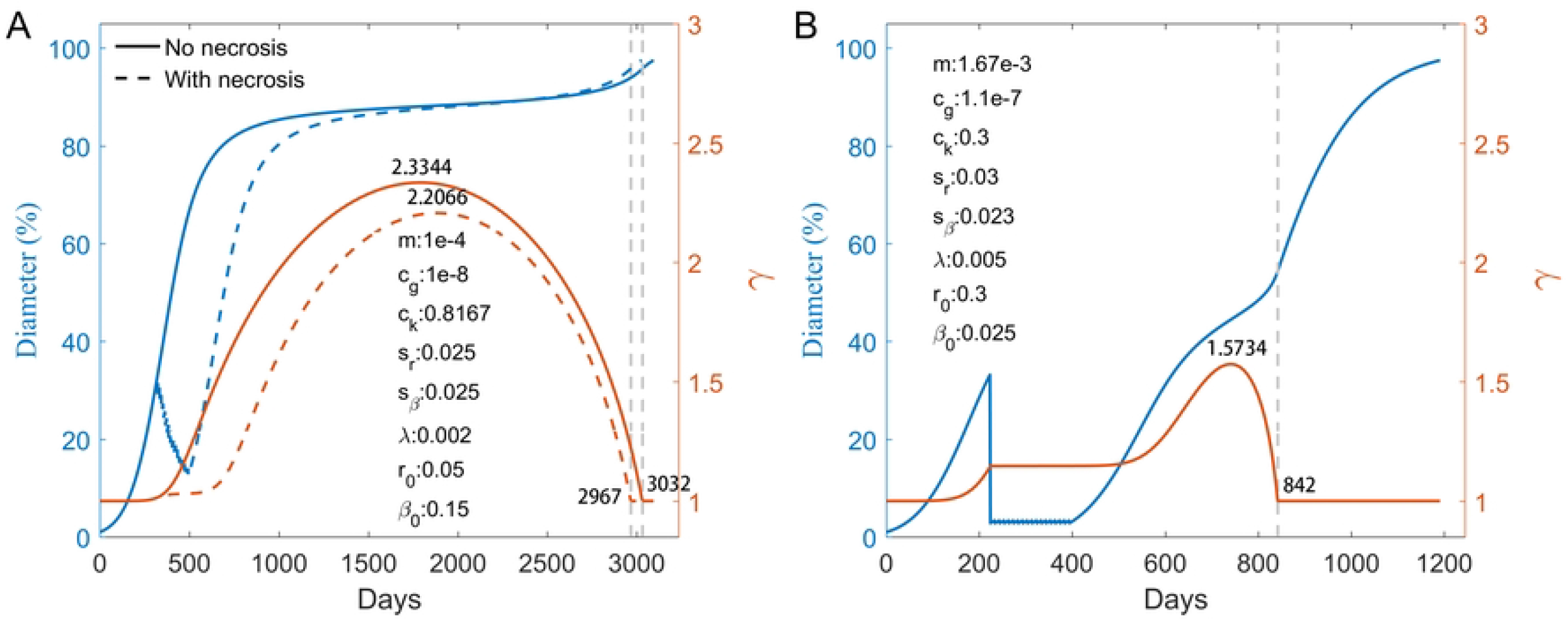
Simulation of necrosis Necrosis was modeled by cell death in proportion to the r to β ratio of each clone that occurred every 10 days for 18 times (Materials and Methods). A. Compared to natural growth without necrosis (blue curve), simulation of necrosis (brown curve) led to shorter overall survival defined by time to 97.5% of K (3584 days and 3501 days, respectively). Peak γ value was lower in necrosis model indicating more invasive behavior as reported ^59^. B. Compared to natural growth demonstrated in Figure 3A and Figure S1A, necrotic model again showed shorter overall survival (1191 days versus 1318 days) and lower peak γ value (1.573 versus 2.3084).

## Discussion

The r, β and γ setting was analogous to the “three strikes” theory of carcinogenesis, which reminded us that both genetic alterations (mainly represented by r and β) and continuous processes (γ) are involved in tumor progression ^57^. Factors influence tumor growth in an overlapping fashion. For example, vascular structures, cell migration and host immunological status could all affect probability of vascular invasion and metastases ^1, 3, 4, 8, 9^. It would be unrealistic to represent each factor categorically in a model of global cancer growth. Parameters in our model were therefore phenomenological, and represent the composite effect of different factors ^2, 10, 16, 17, 20–31^.

K and γ represented environmental restrictions on growth of all cancer cells. K stood for the ultimate capacity of human body. There are other factors that would potentially stall proliferation in a global way though, which are collectively represented by γ. Physical restraint by tumor capsule, pseudo-capsule or external compression is the main source of γ type constraint at early stage ^51^. It would explain sigmoidal growth of primary tumors. Being malignant, such restriction would gradually be overcome by local invasion and metastasis that were assumed to happen stochastically ^3, 9, 12, 38–40^. We have accordingly set the γ curve (equation 2) to gradually decrease after increasing at the beginning. Increase of total number of cancer cells would then start to accelerate, with a full release when γ came back to 1 (or it could come back to any other number, which calls for future investigation), corresponding to tumor progression at advanced stage during which local influences are minimal and systemic factors such as immune responses are the most prominent pathophysiological constituent of γ type restrictions. For blood cancers like CLL, γ should therefore be changing minimally and mutation rate would be the main influencer of growth curve, echoing the reported coexistence of exponential, logistic and intermediate growth in this type of cancer and their relation to mutational traits ^14^. (Figures 2) Notably, this coexistence of several standard types of growth in one kind of cancer is obviously not allowed by any one of the standard models, and this strong effect of mutation when there is no physical boundary to limit tumor growth has also been demonstrated by other simulations ^1, 37^. In these studies, exponential or super-exponential growth was driven by mutation and more malignant colonies replacing older and less aggressive ones, a process that required when cell density is considered, Neumann boundary conditions ^1, 37, 38^, exactly as in our model (See also Figure S3E). In our model however free movement across edge of tumor is not a feature that homogenously started at tumorigenesis^1^ but more of a percentage that increased as cross-border movement happened more often and microsatellite lesions and distant metastases accumulated, which is reflected by the γ curve. We have to hereby point out a weakness of phenomenological description that is the lack of precision. Currently evidence for analysis of the proportional contribution of invasive growth to the increment of total cell number in relation to expansive growth of the primary lesion and metastatic growth at advanced stage is insufficient for clear-cut differentiation. Medical imaging may designate growth of local invasion as part of enlargement of primary tumor^1^, thereby explaining the above-mentioned varying growth speed of the primary tumor which is challenging for standard models. Evolving source of observable enlargement of the primary lesion would also account for clinically observed independence of the spread of the disease from the size of the index tumor ^2, 18, 19, 55^.

Differences between our model and standard models in mathematical interpretation of primary, secondary, and global tumor growth would lead to divergence in how we read into clinical data for prediction of inception time of metastases. Iwata, K. et al first developed the method built on standard sigmoidal growth theory. To account for clinically undiscernible newly started lesions they used a colony size distribution variable *ρ*(*x,t*) which multiplying a cell number Δ*x* indicated the number of colonies that contained *x* to *x* + Δ*x* cells at time *t*. Then assuming all colonies of size *x’* would grow at *g*(*x’*), and that metastases originated both from the primary lesion *x_p_* (primary seeding) and from metastatic lesions (secondary seeding), the authors found an explicit solution for *ρ*(*x,t*) and solved for all parameters based on consecutive clinical measurement of both primary and metastatic lesions of a liver cancer patient. Parameter *b* which represented tumor size at saturation level was thus predicted to be 7.1 × 10^10^ cells which according to the set-up of the report would amount to a tumor of which the diameter was 5 cm. This is far less than the size a real-world liver cancer can grow to ^19^. Our model would justify such incongruency by defining some of the metastatic lesions to be local satellite nodules that on clinical images would be measured as integral parts of primary lesion^1^. However, for these models based on standard equations, this would significantly complicate the subsequent process of deduction for probable inception time of first metastasis. Because the assumption that all lesions grew according to one Gompertzian equation and that all metastatic lesions are distant was fundamental for most studies built on this mathematical framework ^12, 39, 40^. Inserting other reported values into equation 6 would generate prediction of size of the largest metastatic colony 559 days or 632 days after the first diagnosis of the primary tumor quite different from shown in Figure 4a of the report ^40^. The assumption of all colonies with the same number of cells grew at the same speed was questionable too. And although the uniform Gompertzian growth law was simple enough, calculating the summed effect of primary and secondary dissemination was complicated to such an extent that other studies have been considering only primary dissemination to avoid confusion ^12, 39^. However as shown above, size prediction of primary lesion by such method led to significant underestimation, let alone representing all lesions by it. Agreeing with the general idea, our model would differ by providing a gross prediction of all surviving cells including the source of both primary and secondary seeding. Or to simplify computational process, we would recommend exponential instead of standard sigmoidal equation to represent growth of all cancer cells instead of those in primary lesion. The computational process is much simpler and assuming much less than the colony size distribution method. Starting from clinically observed data, identifiability is almost guaranteed without significant inconsistency. Starting from size measurements of the largest metastasis, we predicted that the first distant metastasis started well after diagnosis of the primary lesion, which was much later than the predicted time in the original report. This was a difference not unworthy of attention and further investigation.

A particular theory in HCC patients further illustrates independent prognostic value of lesion multiplicity. Among multinodular HCCs, multicentric occurrence (MO) type was claimed to exhibit significantly less malignant features and longer survival after surgery than intrahepatic metastasis (IM) type ^56^ with primary lesions of similar sizes. This phenomenon would be inexplicable in standard models since dissemination rate in these models related directly to cell number while proliferation advantage parameter in standard equations were set to distribute evenly ^12, 38–40^. In our model however, MO type could be described by smaller C_k_, smaller C_γ_ and diagnosis before γ returned to 1. IM type indicated the point of growth where γ restriction had been breached (Figure1a). IM type therefore should contain more undetected metastases, constitute a larger proportion of clones generated later, and grow faster at this point (Figures 1a-b and see also Figure S2), agreeing to the reported phenomenon ^56^. Others have also proposed independence of invasion and metastasis from index tumor, stating that “tumor growth rate became rapid at the time of recurrence” ^18^ which was also observed in our simulation for IM and MO type tumors (Figure 1).

Recently, coupling theory predicted increased heterogeneity in exponential growth, which was the exact opposite of the sweeping model previously predicted for exponential growth^1, 5, 37^. The inconsistency originated from different definitions of heterogeneity. It was represented in the coupling theory by parameters m and f(n) that described the availability of free states^5^. In our model and standard exponential models describing sweeping mechanism, heterogeneity was represented by not m but parameters *a* and α in the coupling model that denoted respectively growth rate and the proportion of proliferative states among all possible states^1, 5^. Freedom in our model was represented by γ. This divergence demonstrated the use of a model properly describing global cancer growth where multiple parameters are reserved to classify and represent the complicated networks of biological processes involved to avoid confusion. Meanwhile, instead of sweeping mode or big bang theory ^1, 50^ we made more complicated prediction of heterogeneity (See also Figure S1-3): although simulation indicated that the earliest clones tended to be swept, they could maintain dominance when mutation rate was low (See also Figure S3D), or clones starting from slightly later would (See also Figure S1C&D), because over a certain point the 1-(N(t)/K) limitation could become too severe for new clones, while earlier clones would keep growing because of bigger population base (See also Figure S1-3). Nonetheless compared with early stage, release from γ type boundary restriction at advanced stage would result in faster evolution driven by more aggressive clones generated later in time and concurrently faster (explosive) growth of the total cancer population, as predicted by previous reports (See also Figure S1C&D) ^1, 37, 38^. Therefore our model is not refusing any of the hypotheses of heterogeneity. Instead, we think it predicts a spectrum of possibilities that could contain explanatory discrepancies when variations in mutation mechanism, distribution of γ, sampling error related to biopsy, and interpretation of sequencing results were all considered ^1, 5, 13, 14, 53^. More importantly its prediction of the relation between evolution, regional restriction and growth speed coincided with previous reports ^1, 37, 38^.

Heterogeneity would also be preserved during our treatment simulation, as predicted by the DTP theory (Figure 3) ^58^. Three main features of our model suggested that it would be well fitted for simulating treatment at advanced stage. Firstly, it considers all cancer cells from all lesions at this stage simultaneously. Secondly, γ at advanced stage represents systemic factors stalling cancer growth which are practically the roles played by systemic antitumor medications. Increase in γ would affect all cancer cells equally and simultaneously which reflected the conservative feature of DTP state. Interestingly, our model also supported the DTP theory by illustrating that if treatment reduced cell number without additional suppression from increased γ, it would lead to less changes in prognosis (Figure 3 and see also Figure S4). It was also depicted that if given at advanced stages, even treatment that induced an arbitrary increase of γ would need to be repeated to prolong survival, which echoed clinical routine (Figure 3 and see also Figure S4). And thirdly parameter K described carrying capacity of a patient, which would be convenient for simulation of side-effect of systemic treatment (not covered in the present study). Noticeably, in an earlier modelling study based on clinical observation, it was suggested that global reduction of tumor growth rate was more efficient in extending patient life expectancy than physical removal of primary lesion. The authors stated that “Surgical resection of the primary tumor, even if done efficiently such that 99.99% of the primary tumor was removed, led to less promising results” ^2^. This was almost exactly identical to our treatment simulation results (Figure 3 and see also Figure S4). Potentiality of our model to approximate systemic anti-tumor treatment in a realistic and detailed fashion is meaningful not only in validation of its identifiability. Improved modeling of treatment effects in the future may help reduce cost of clinical trials, especially the need for large number of participants.

Necrosis simulation was also performed as an independent identifiability test ^59^. Again, the reported observation that although causing additional cell loss, necrosis was related with shorter survival and more invasive behavior was impossible for standard models, but readily simulated by our model (Figure 4).

However, these simulations were only preliminary. Much refinement needs to be done in the future. A major disadvantage of our model was that growth of tumor volume was considered linearly correlated to number of viable tumor cells, excluding effect of compression and other components of a cancer lesion. Although general oncological theories seemed to be not suggesting otherwise, further investigation is needed.

### Conclusion

This inhomogeneous generalized model is compatible with distinct clinical observations that are not well explained by existing modelling theories including varying growth speed of primary lesion, coexistence of exponential, sigmoidal and intermediate growth in leukemia, independent prognostic value of multiplicity of lesions, contradicting discoveries of heterogeneity evolution, treatment effect and necrosis related clinical behavior. It provided a phenomenological perspective for explaining clinical observations by mathematical model that was different from mechanism-based deterministic perspective in other modelling studies. Since both have exhibited acceptable identifiability, it would be interesting for future investigation to compare the two methods. If we look at modeling studies as trying to link laboratory data to clinical observation, the current model indicated that the bridging work could be done along the opposite direction.

## Author Contributions

H. W., Y. M., Z. Z., L. L., W. L., and M. H. designed the study, analysed and interpreted data. H. W., Y. M. and B. C. modeled tumor growth. H. W., S. L., S. Z., B. C., and H. Wen. conducted extensive search through published documents and analysed goals of simulation. Y. M., Z. Z., Z. X., and W. L. developed the code for simulations. Z. X., and W. L. made analytic calculations. All authors discussed the results. The manuscript was written primarily by H. W., Y. M., L. L., W. L., and M. H., with contributions from Z. Z. All authors read and approved the final manuscript.

## Acknowledgments

We are grateful to Suohai Fan, and Weidong Wang for helpful discussions.

## Declaration of Interests

The authors declare no potential conflicts of interest.

## Notes

### Competing Interest Statement

The authors have declared no competing interest.

